# Recombinant protein subunit SARS-CoV-2 vaccines formulated with CoVaccine HT adjuvant induce broad, Th1 biased, humoral and cellular immune responses in mice

**DOI:** 10.1101/2021.03.02.433614

**Authors:** Chih-Yun Lai, Albert To, Teri Ann S. Wong, Michael M. Lieberman, David E. Clements, James T. Senda, Aquena H. Ball, Laurent Pessaint, Hanne Andersen, Oreola Donini, Axel T. Lehrer

## Abstract

The speed at which several COVID-19 vaccines went from conception to receiving FDA and EMA approval for emergency use is an achievement unrivaled in the history of vaccine development. Mass vaccination efforts using the highly effective vaccines are currently underway to generate sufficient herd immunity and reduce transmission of the SARS-CoV-2 virus. Despite the most advanced vaccine technology, global recipient coverage, especially in resource-poor areas remains a challenge as genetic drift in naïve population pockets threatens overall vaccine efficacy. In this study, we described the production of insect-cell expressed SARS-CoV-2 spike protein ectodomain and examined its immunogenicity in mice. We demonstrated that, when formulated with CoVaccine HT™adjuvant, an oil-in-water nanoemulsion compatible with lyophilization, our vaccine candidates elicit a broad-spectrum IgG response, high neutralizing antibody titers, and a robust, antigen-specific IFN-γ secreting response from immune splenocytes in outbred mice. Our findings lay the foundation for the development of a dry-thermostabilized vaccine that is deployable without refrigeration.

## Introduction

The emergence and rapid spread of a new infectious respiratory disease, Coronavirus Disease 2019 (COVID-19) has caused an unprecedented public health emergency worldwide since emerging in December 2019. A novel coronavirus, severe acute respiratory syndrome coronavirus 2 (SARS-CoV-2), which is closely related to SARS-CoV, was identified as the etiologic agent of the new respiratory disease. Despite the practice of public health strategies such as social distancing and use of face masks, numbers of COVID-19 cases continue to rise globally. As of March 2021, there are over 114 million confirmed cases with over 2.4 million deaths worldwide and over 28 million cases and 513,000 fatalities reported in the U.S. [1], accentuating the urgent need for safe and effective, inexpensive and easily deployed vaccines that would rapidly establish herd immunity to break transmission on a global scale. Facilitated by the broad nature of the outbreak, significant SARS-CoV-2 variants have been identified, some of which appear to be more resistant to first-generation vaccines. More than 200 COVID-19 vaccine candidates using various technology platforms are currently in development [2]. Among these, two frontrunning vaccines based on mRNA platforms, Pfizer/BioNTech BNT162b2 and Moderna mRNA-1273 with reported interim efficacy rates of 95% and 94.1% respectively, have been approved by the U.S. Food and Drug Administration (FDA) for emergency use in mid-December 2020 [3, 4]. These two vaccines are now being administered in the U.S. and have also been approved for use in other jurisdictions with most authorities recommending administration to high-risk individuals only as supply is limited. Furthermore, both vaccines require stringent cold-chain distribution and storage. For the furthest advanced vaccines at least one booster dose seems necessary, bringing global demand to at least 16 billion doses [5]. While the recent Emergency Use Authorization (EUA) approval of the Janssen Ad26.COV2.S vaccine, which showed 85% efficacy in preventing severe COVID-19, is expected to ease this burden [6, 7], no single vaccine can be produced rapidly in sufficient quantities to satisfy the global urgent need, diversification using different vaccine platforms would enable worldwide vaccine coverage as well as address the needs of the most vulnerable populations, particularly elderly and immunocompromised individuals or those with other co-morbidities. With mutations emerging even in the absence of significant selection pressure placed on SARS-CoV-2, the initiation of vaccination programs may further enhance strain diversity. Thus, continued research into adaptable and more easily distributed vaccines, compatible with rapid deployment and significant cost efficiencies, must continue unabated.

The recombinant subunit vaccine platform offers a safety advantage over virally vectored vaccines and a distribution advantage relative to many other vaccine platforms. Purified recombinant protein antigens can be engineered to achieve optimal immunogenicity and protective efficacy. Furthermore, a thermostabilized subunit vaccine can be deployed in the field, eliminating stringent cold-chain requirements. Formulation of the vaccine immunogen with a potent adjuvant enhances and focuses immunogenicity while lowering the antigen dose requirement, thereby enabling vaccination of more people with a product carrying significantly more clinical and regulatory precedence compared to nucleic acid-based approaches.

CoVaccine HT™, an oil-in-water nanoemulsion adjuvant with excellent safety, immunogenicity and stability, in combination with properly selected antigens can achieve potent immunogenicity and protective efficacy in rodents and non-human primates (NHPs) [8]. In previous work, we have successfully demonstrated the use of recombinant protein subunit *Drosophila* S2 expression system in combination with CoVaccine HT™ to produce vaccines to combat global health threats such as Zika virus (ZIKV) and Ebola virus (EBOV). Immunization with recombinant ZIKV E protein induced potent neutralizing titers in mice [9] and non-human primates [8] and protection against viremia after viral challenge. Similarly, immunization with recombinant subunit formulations consisting of the EBOV glycoprotein and matrix proteins VP40 and VP24 was able to induce potent antibody titers and protection in both mouse [10] and guinea pig models [11]. More recently, we have demonstrated that these recombinant subunits, in combination with CoVaccine HT™, can be formulated as a glassy solid using lyophilization. In this format, lyophilized vaccine formulations consisting of filovirus glycoprotein and adjuvant were able to maintain native quaternary structure and immunogenicity for at least 12 weeks at 40 °C [12, 13]. Other vaccines, using a similar lyophilization process, demonstrated potency after at least 1 year at 40 °C [14]. Thus, we have the capacity to produce a lyophilized, adjuvanted recombinant protein vaccine in a single-vial, which is stable for long term storage even at elevated temperatures and which can be reconstituted with sterile water for injection immediately prior to use. Spike (S) glycoprotein, comprised of a receptor binding subunit (S1) and a membrane-fusing subunit (S2) [15] is the main surface protein and present as homotrimers on the viral envelope of SARS-CoV-2. Based on previous preclinical studies of vaccines against the highly pathogenic SARS-CoV and Middle East respiratory syndrome coronavirus (MERS-CoV) [16–18] as well as recent studies of patients with SARS-CoV-2 infections [19–22], S protein appears to be the antigenic target of both neutralizing antibody and T cell responses. The majority of current COVID-19 vaccines under preclinical and clinical development use full-length S proteins as antigen targets with further modifications such as removal of the polybasic sites [23–25], introduction of proline mutations [23, 26, 27], or addition of trimerization domains to preserve the native-like trimeric prefusion structure of S proteins. These antigens have been shown to mimic the native S protein presented on viral particles and preserve neutralization-sensitive epitopes [18, 28]. In a prior study, we evaluated the utility of CoVaccine HT™ adjuvant to induce properly balanced immunity against SARS-CoV-2, when formulated with a commercially available SARS-CoV-2 spike S1 protein. This work demonstrated that CoVaccine HT™ is an effective adjuvant that promotes rapid induction of balanced humoral and cellular immune responses [29]. In the current study we produced a native-like trimeric S protein ectodomain with and without stabilizing mutations using the *Drosophila* S2 cell expression system and assessed the immunogenicity of these S ectodoman trimers formulated with CoVaccine HT™ in mice.

## Materials and Methods

### 2.1 Ethics Statement

All animal work was conducted in accordance with the Animal Welfare Act and the National Research Council (NRC) Guide for the Care and Use of Laboratory Animals. All experimental procedures were reviewed and approved by the Institutional Animal Care and Use Committee (IACUC) at the University of Hawaii at Manoa (UHM) and carried out in the UHM American Association for Accreditation of Laboratory Animal Care (AAALAC) accredited Laboratory Animal Facility.

### 2.2 Recombinant protein expression and purification

Plasmids were generated to express the native-like, trimeric, transmembrane (TM)-deleted spike (S) glycoprotein (SdTM) from SARS-CoV-2 strain Wuhan-Hu-1 (Genbank Accession number NC_045512). The SdTM sequence was designed to encode the SARS-CoV-2 S protein sequence spanning Gln14 to Ser1147. The SdTM gene was produced by de novo synthesis (ATUM, Newark, CA). The gene was also codon-optimized for expression in *Drosophila* S2 cells, with an altered furin cleavage site (RRAR changed to GSAR) between S1 and S2 domains to prevent cleavage, and contains a trimerization domain of T4 bacteriophage fibritin (foldon) at the C-terminus. Two additional proline substitutions (K986P and V987P) between the heptad repeat 1 and central helix regions and the removal of the S2’ protease cleavage site were introduced by site-directed mutagenesis to generate the stabilized prefusion structure of S protein (SdTM2P). SARS-CoV-2 receptor binding domain with foldon trimerization domain at the C-terminus (RBD-F) was also prepared. The RBD-F sequence was designed to encode the SARS-CoV-2 S protein sequence spanning Phe318 to Gly594. The proprietary expression vector pHH202 (Hawaii Biotech Inc., Honolulu, HI) containing S gene variants were transfected into *Drosophila* S2 cells using the Lipofectamine LTX with PLUS reagent (Invitrogen, Carlsbad, CA) according to manufacturer’s instructions. Stably transformed cell lines were created by selection with culture medium containing hygromycin B at 300 μg/mL. To verify selection, transformants were induced with culture medium containing 200 μM CuSO4. Expression was verified by sodium dodecyl sulfate polyacrylamide gel electrophoresis (SDS-PAGE) and western blotting.

Recombinant S proteins were purified from filtered cell culture supernatants by affinity chromatography (AC) using NHS-activated Sepharose (Cytiva, Marlborough, MA) coupled with 2 mg of a his-tagged human angiotensin I converting enzyme 2 (hACE2) produced using the *Drosophila* S2 cell expression system and Ni-affinity chromatography in our laboratory. Purified recombinant S proteins were concentrated using Amicon filtration devices (EMD Millipore, Billerica, MA), buffer-exchanged into PBS and analyzed by SDS-PAGE and Western blotting. Antigens were quantified by UV absorbance at 280 nm and stored at −80°C. A conventional immunoaffinity chromatography (IAC) method was also established to purify SdTM2P proteins. For this, the monoclonal antibody (mAb) CR3022 (provided by Mapp Biopharmaceutical) was coupled to NHS-activated Sepharose at a concentration of 10 mg/mL and used for IAC in tandem with a HiPrep 26/10 Desalting column (Cytiva, Marlborough, MA) equilibrated with PBS allowing quick buffer exchange of the eluted protein from low pH buffer into PBS.

### 2.3 Mouse experiments

Groups (n=7 or 15 per group) of 7 to 10-week-old Swiss Webster mice of both sexes (bred from original breeding stocks obtained from Taconic Biosciences, Germantown, NY) were immunized intramuscularly (i.m.) at days 0 and 21 with 5 μg of SARS-CoV-2 spike proteins SdTM or SdTM2P (purified by hACE2 affinity chromatography) alone or in combination with 1 mg of CoVaccine HT™ adjuvant (Protherics Medicines Development Ltd, a BTG company, London, United Kingdom). The negative control group received equivalent doses of adjuvant only. To determine the optimal antigen and adjuvant dosing, mice were also immunized with either 5, 2.5, or 1.25 μg of SdTM2P (purified by mAb CR3022 IAC) and 1 mg of CoVaccine HT™ or 5 μg of SdTM2P formulated with 0.3 mg CoVaccine HT™. Serum samples were collected on days 7, 14, 28, and 35. Three to five mice from each group were euthanized on the seventh day after the first and second vaccinations and splenectomies were performed for preparation of splenocytes. The remaining four to five mice from each group were euthanized 14 days after the second vaccination and terminal bleeds collected by cardiac puncture.

### 2.4 Analysis of antibodies by multiplex microsphere immunoassay (MIA)

The IgG antibody in mouse sera was measured by a multiplex microsphere-based immunoassay as described previously [9, 29, 30]. Briefly, internally dyed, magnetic MagPlex® microspheres (Luminex Corporation, Austin, TX) were coupled to purified receptor binding domain (RBD-F), spike protein (SdTM2P) or bovine serum albumin (BSA) as control [9, 29]. A mixture of RBD, spike SdTM2P, and BSA-coupled beads (approximately 1,250 beads each) was incubated with diluted sera in black-sided 96-well plates for 3 hours at 37°C with gentle agitation in the dark. Following two washes with MIA buffer (1% BSA and 0.02 % Tween 20 in 1x PBS), 50μl of 1μg/mL red phycoerythrin (R-PE)-conjugated goat anti-mouse IgG antibodies (Jackson ImmunoResearch, Inc., West Grove, PA) were added and incubated at 37°C for another 1 hour. After washing twice with the MIA buffer, the beads were resuspended in MAGPIX® drive fluid and analyzed on a MAGPIX® Instrument (Luminex Corporation, Austin, TX).

The median fluorescence intensity (MFI) readouts of the experimental samples were converted to antibody concentrations using purified antibody standards prepared from pooled mouse antiserum to SdTM or SdTM2P as described below. For this, IgG was purified from mouse antisera by protein A affinity chromatography and then subjected to immunoaffinity chromatography (IAC) using SdTM2P-coupled NHS-Sepharose (Cytiva, Marlborough, MA) to select only S-reactive IgG. The concentration of purified S-specific IgG was quantified by measuring the UV absorbance of the solution at 280 nm. The purified anti-spike IgG was diluted to concentrations in the range of 4.88 to 5000 ng/mL and analyzed in the MIA assay. The resulting MFI values were analyzed using a sigmoidal dose-response, variable slope model (GraphPad Prism, San Diego, CA), with antibody concentrations transformed to log10 values. The resulting curves yielded r^2^values > 0.99 with well-defined top and bottom and the linear range of the curve was determined. The experimental samples were analyzed side-by-side with the antibody standards at different dilutions (1:50, 1:200, 1:1000, 1:40,000, 1:80,000 or 1:160,000) to obtain MFI values that fall within the linear range of the standard curve. The experimental sample IgG concentrations were interpolated from the standard curves using the same computer program. Finally, the interpolated values were multiplied by the dilution factors and plotted as antibody concentrations (ng/mL).

The IgG subclass profile in serum samples was analyzed using IgG subclass-specific secondary antibodies (Southern Biotech, Birmingham, AL), and the ratios of IgG2a/IgG1 and IgG2b/IgG1 were calculated using the MFI readouts at the serum dilution (1:2000) that is within the linear range of the antibody binding standard curve.

### 2.5 Recombinant vesicular stomatitis virus (rVSV) plaque and neutralization assay

Replication-competent rVSV expressing SARS-CoV-2 S protein without cytoplasmic tail (rVSV-SARS-CoV-2-S) was provided by Dr. Andrea Marzi (Laboratory of Virology, National Institute of Allergy and Infectious Diseases) and the virus stocks were amplified in Vero E6 cells. Virus titers were measured on Vero E6 cells in 6-well plates by standard plaque assay. Briefly, 500 μL of serial 10-fold virus dilutions were incubated with 2.5x 10^6^cells/well at 37°C for 1 hour and then overlaid with DMEM containing 2% fetal bovine serum (FBS) and 1% agarose. Following incubation in a 37°C, 5% CO_2_ incubator for 72 hours, cells were fixed and stained with a solution containing 1% formaldehyde, 1% methanol, and 0.05% crystal violet overnight for plaque enumeration. For the plaque reduction neutralization test (PRNT) using rVSV-SARS-CoV-2-S, pooled or individual mouse serum samples were heat-inactivated at 56°C for 30 minutes. Eight 3-fold serial dilutions of serum samples starting at final 1:10 dilution were prepared and incubated with 100 plaque-forming units (PFU) of rVSV-SARS-CoV-2-S at 37°C for 1 hour. Antibody-virus complexes were added to Vero E6 cell monolayers in 6-well plates and incubated at 37°C for another hour followed by addition of overlay media. Three days later, the plaque visualization and enumeration steps were carried out as described in the plaque assay. The neutralization titers (PRNT_50_) were defined as the highest serum dilution that resulted in 50% reduction in the number of plaques.

### 2.6 Wild-type SARS-CoV-2 virus neutralization assay

PRNT was also performed in a biosafety level 3 facility at BIOQUAL, Inc. (Rockville, MD) using 24-well plates. Mouse sera pooled from individual mice within each group, were diluted to 1:10, and a 1:3 serial dilution series was performed 11 times. Diluted samples were then incubated with 30 plaque-forming units of wild-type SARS-CoV-2 (USA-WA1/2020, BEI Resources NR-52281) in an equal volume of culture medium (DMEM with 10% FBS and gentamicin) for 1 hour at 37°C. The serum-virus mixtures were added to a monolayer of confluent Vero E6 cells and incubated for 1 hour at 37°C in 5% CO_2_. Each well was then overlaid with 1 ml of culture medium containing 0.5% methylcellulose and incubated for 3 days at 37°C in 5% CO_2_. The plates were then fixed with methanol at ‒20°C for 30 minutes and stained with 0.2% crystal violet for 30 min at room temperature. Neutralization titers (PRNT_50_) were defined as the highest final serum dilution that resulted in 50% reduction in the number of plaques.

### 2.7 Splenocyte preparation and FluoroSpot

Three or five mouse spleens from each group were harvested seven days after the first and second vaccinations, and single cell suspensions were prepared using a gentleMACS Dissociator (Miltenyi Biotec, Auburn, CA). The cells were passed through a cell strainer, resuspended in freezing medium containing 90% FBS and 10% dimethyl sulfoxide (DMSO) after lysis of red blood cells, and cryopreserved in liquid nitrogen. FluoroSpot assay was performed using mouse IFN-γ FluoroSpot^PLUS^kit according to the manufacturer’s instructions (Mabtech, Inc., Cincinnati, OH). Briefly, splenocytes were rested in a 37°C, 5% CO_2_ incubator for 3 hours after rapidly thawing in a 37°C water bath followed by slow dilution with culture medium to allow removal of cell debris. A total of 2.5 × 10^5^cells per well in RPMI-1640 medium supplemented with 10% FBS, penicillin (100 units/mL) and streptomycin (100 μg/mL) were added in a 96 well PVDF membrane plate pre-coated with capture monoclonal antibodies. The cells were stimulated for 24 hours with a peptide pool consisting 17-mer peptides with 10 amino acids overlapping, covering the spike protein of SARS-CoV-2 (BEI Resources NR-52402) at 10 μg/mL or medium containing equal concentration of DMSO (0.05%) as negative control. Fifty thousand cells were incubated with cell activation cocktail (BioLegend, San Diego, CA) at 1:500 dilution containing phorbol-12-myristate 13-acetate (PMA, 0.081 μM) and Ionomycin (1.3386 μM) as positive control. Each stimulation condition was set up in triplicate. Plates were developed using specific monoclonal detection antibodies and fluorophore-conjugated secondary reagents. Finally, plates were treated with a Fluorescence enhancer (Mabtech) to optimize detection and then air-dried. The spots were enumerated using the CTL ImmunoSpot® S6 Universal Analyzer (Cellular Technology Limited, CTL, Shaker Heights, OH), and the number of antigen-specific cytokine secreting spot forming cells (SFCs) per million cells for each stimulation condition was calculated by subtracting the number of spots detected in the medium only wells.

### 2.8 Statistical analysis

Statistical analysis was performed using one sample t test or Mann-Whitney t test to compare the neutralization titers, IgG subclass antibody profile, and IFN-γ secreting response between adjuvanted and unadjuvanted groups. The difference in the neutralizing antibody titers and the numbers of IFN-γ secreting cells between groups receiving different dosages of vaccines was determined by one-way ANOVA with Tukey’s multiple comparison test. The correlation between the measured neutralization titers using rVSV-SARS-CoV-2 and authentic SARS-CoV-2 was analyzed by Pearson correlation analysis.

## Results

### 3.1 Drosophila S2 cell-expressed recombinant S proteins adjuvanted with CoVaccine HT™ induce robust IgG antibody responses

Using the *Drosophila* S2 cell expression system, we generated trimeric SARS-CoV-2 S protein devoid of transmembrane domain (SdTM) as well as a stabilized prefusion structure of S protein (SdTM2P). To explore different methods for affinity purification of S protein, we first employed a novel method in which we utilized human angiotensin I converting enzyme 2 (hACE2) protein, a known receptor for SARS-CoV-2, for AC purification of S proteins. The immunogenicity of recombinant S proteins was evaluated with and without CoVaccine HT™ adjuvant in outbred Swiss Webster mice. Animals were immunized with one or two doses at a 3-week interval by intramuscular injection with 5 μg of SARS-CoV-2 SdTM or SdTM2P (purified by hACE2 AC) alone or formulated with 1 mg CoVaccine HT™ (Fig. 1A). We developed a quantitative IgG assay in which purified anti-S polyclonal antibody was used as a standard and therefore the antigen-specific IgG levels are expressed as concentration (ng/mL) (Fig. 1B). Analysis of IgG antibody in sera obtained on days 7, 14, 28, and 35 indicated that the co-formulation of CoVaccine HT™ with SdTM or SdTM2P significantly increased anti-S IgG antibody levels over those induced by proteins alone (Fig. 1C). In addition, two doses of CoVaccine HT™ adjuvanted SdTM or SdTM2P elicited high levels of IgG antibodies to RBD, which is the target of neutralizing antibodies (Fig. 1D). To further optimize vaccine formulations, we evaluated the IgG antibody responses in mice that received a lower dosage (2.5 or 1.25 μg) of SdTM2P proteins or that received a reduced amount (0.3 mg) of adjuvant. The results demonstrated that vaccination with reduced amounts of antigen or adjuvant did not decrease the levels of antigen-specific IgG antibodies (Fig. 1E).

**Fig. 1.**
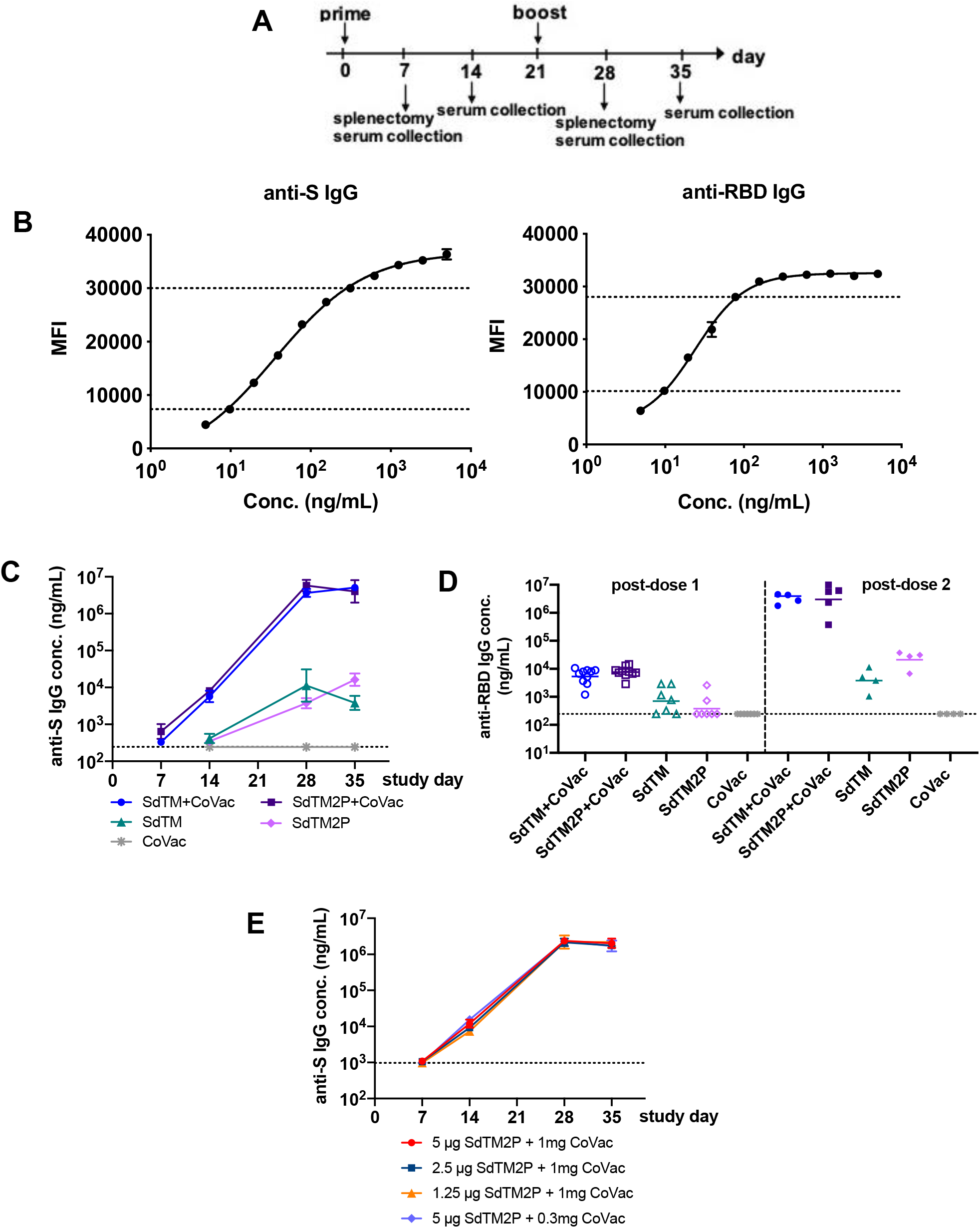
IgG antibody responses to recombinant SARS-CoV-2 S proteins. (A) Groups of Swiss Webster mice (n=7 or 15) were immunized with one or two doses of recombinant S proteins with or without CoVaccine HT™ (CoVac) adjuvant at a 3-week interval. Sera were collected 1 and 2 weeks after each immunization (days 7, 14, 28, and 35), and spleens were harvested 1 week (day 7 and 28) after each immunization. SARS-CoV-2 S-specific IgG titers were measured by a multiplex microsphere immunoassay (MIA) using SdTM2P and RBD-F coupled beads. (B) The purified anti-S antibody was diluted to concentrations in the range of 4.8 to 5000 ng/mL and analyzed by MIA as a standard (as described in the Materials and Methods). Mouse sera were assayed along with the antibody standard and the IgG concentrations was interpolated from the standard curves using a sigmoidal dose-response computer model (GraphPad Prism). The dotted lines denote the top and bottom of linear range that were used to interpolate antibody concentration (C) The anti-S and (D) anti-RBD antibody titers in sera from mice immunized with SdTM or SdTM2P (purified by hACE2 AC) with or without adjuvants or (E) anti-S antibody in sera of mice administered different dosages (5, 2.5, or 1.25 μ;g) of SdTM2P (purified by mAb IAC) with adjuvant (1 or 0.3 mg) are expressed as IgG concentrations (ng/mL). The dotted lines in panels C to E indicate the bottom of linear range of the standard curve.

### 3.2 Recombinant S proteins adjuvanted with CoVaccine HT™ elicit potent neutralizing antibody responses

We next examined whether our vaccine candidates induced neutralizing antibody responses. Measurement of SARS-CoV-2 neutralizing antibodies requires biosafety level 3 (BSL3) laboratory facilities, and therefore, alternative assays have been developed, such as one utilizing replication-competent rVSV-ΔG expressing SARS-CoV-2 S glycoprotein (rVSV-SARS-CoV-2-S) [31, 32]. We first compared the neutralizing antibody titers of pooled mouse sera using both authentic SARS-CoV-2 and rVSV-SARS-CoV-2-S (Table 1) and found that the titers are significantly correlated between these two assays (Fig. 2C). To further evaluate the neutralizing antibodies of individual animals, we employed only the PRNT using rVSV-SARS-CoV-2-S. The PRNT_50_ titers of sera obtained from mice vaccinated with SdTM or SdTM2P in combination with CoVaccine HT™ were significantly higher than those from animals receiving protein alone (Fig. 2A), suggesting that the CoVaccine HT™ adjuvant enhances induction of neutralizing antibody responses. Furthermore, a trend of dose-dependent decreases of neutralization titers was observed when animals were given lower dosages of SdTM2P proteins (Fig. 2B). Of note, the serum PRNT_50_ titers of mice receiving a lower amount (0.3 mg) of CoVaccine HT™ were comparable or even higher than those of mice given 1 mg of adjuvant indicating that the optimal vaccine formulation may be achieved using lower dosages of adjuvant. The results indicate that as little as 2.5 μg of S protein antigen with 0.3 mg of adjuvant might be sufficient to induce potent neutralizing antibody responses.

**Table 1.**
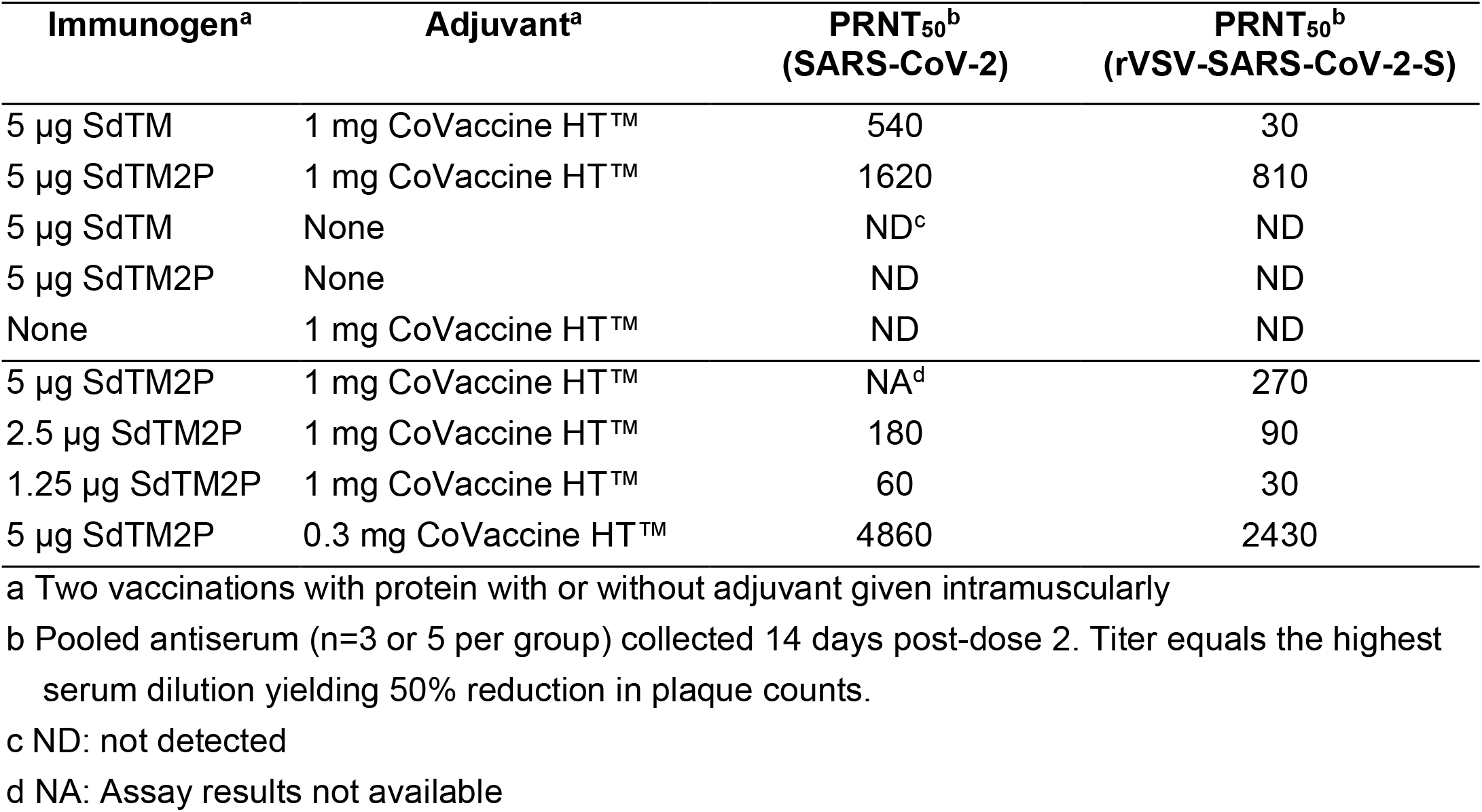
Virus neutralizing antibody titers against SARS-CoV-2 and rVSV-SARS-CoV-2-S in pooled antisera from mice immunized with two doses of S proteins formulated with CoVaccine HT™ adjuvant

**Fig. 2.**
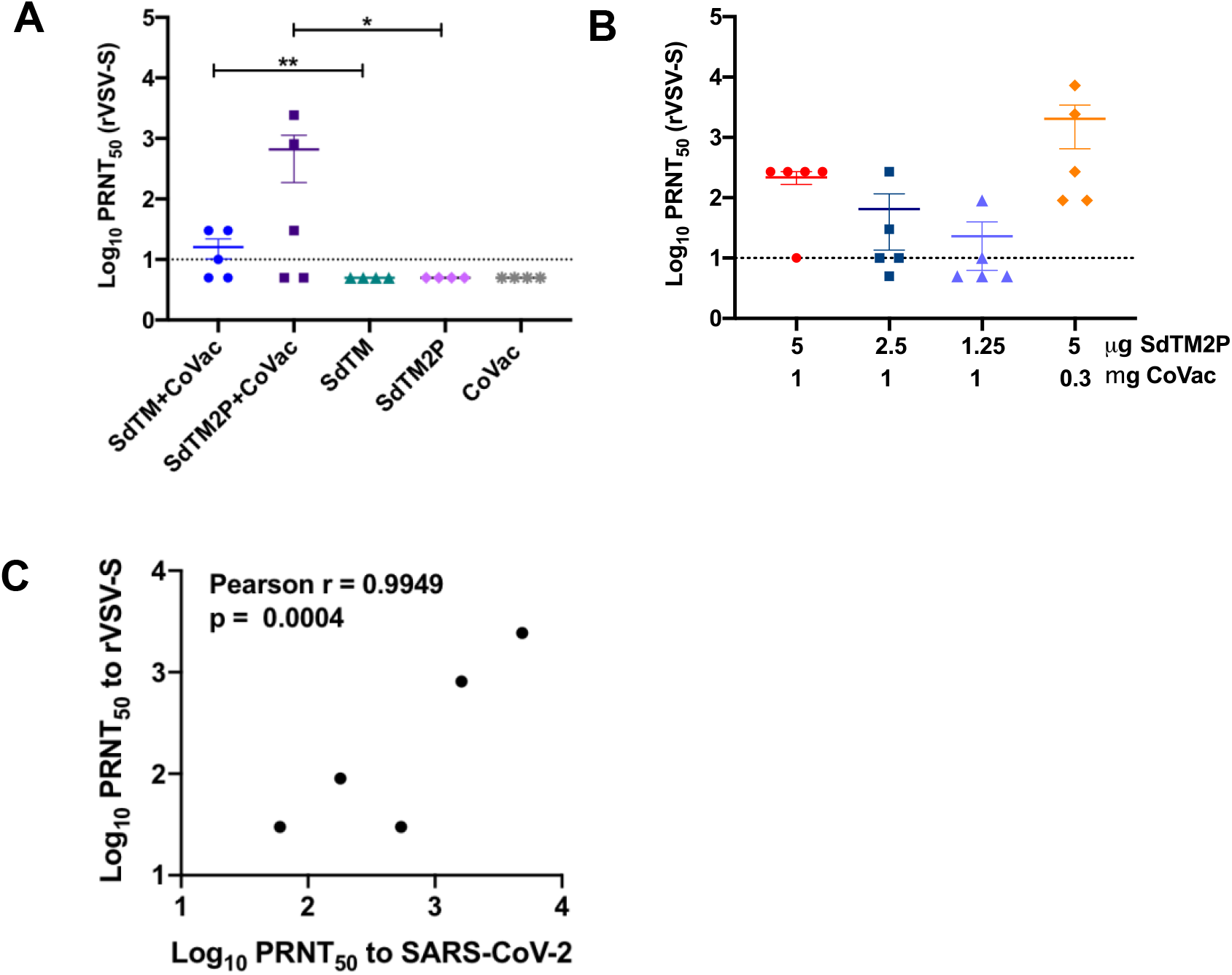
The serum neutralization titers of mice immunized with recombinant subunit SARS-CoV-2 vaccines. The neutralization titers of sera obtained 2 weeks post boosting (day 35) from mice vaccinated with (A) SARS-CoV-2 SdTM or SdTM2P proteins (purified by hACE2 AC) adjuvanted with or without 1 mg of CoVaccine HT™ (CoVac) or (B) different dosages (5, 2.5, or 1.25 μ;g) of SdTM2P (purified by mAb IAC) with CoVac (1 or 0.3 mg) was measured by a PRNT using rVSV-SARS-CoV-2-S. The data shown are log-transformed PRNT_50_ values from individual animals, and mean ± SEM of each group is indicated. Horizontal dashed lines represent the limit of detection. For the samples showing low (<50 %) or no neutralizing activity at the starting dilution (1:10) in the assay, a PRNT_50_ of 5 (half of the limit of detection) was reported when the data were plotted. One sample t test on log-transformed data was used to analyze the significant difference between adjuvanted and protein alone groups in panel A (*p < 0.05, **p < 0.01). There are no significant differences between groups when analyzed by one-way ANOVA with Tukey’s multiple comparisons test in panel B. (C) The correlation between the neutralization titers of pooled serum samples from the same group to rVSV-SARS-CoV-2 and those to WT SARS-SARS-CoV-2 was examined by Pearson correlation analysis. The PRNT_50_ values are listed in Table 1.

### 3.3 Recombinant S proteins in combination with CoVaccine HT™ induce a balanced IgG subtype antibody response

Vaccine-associated enhanced respiratory disease (VARED) has been reported in infants and young children immunized with inactivated whole virus vaccine against respiratory syncytial virus (RSV) and measles virus [33–35], and associated with Th2-biased immune responses [36]. A similar pulmonary immunopathology was also observed in animals with SARS-CoV vaccines [37–39]. Thus we evaluated the balance of Th1 and Th2 by comparing the levels of S-specific IgG2a/b and IgG1, which are indicative of Th1 and Th2 responses, respectively. Both SdTM and SdTM2P adjuvanted with CoVaccine HT™ elicited high anti-S IgG2a/2b and IgG1 subclass antibodies whereas protein alone induced high IgG1 with lower IgG2a and IgG2b (Fig. 3A). To further assess the effect of CoVaccine HT™ adjuvanticity on IgG subclass antibody profiles, the ratios of IgG2a versus IgG1 as well as IgG2b versus IgG1 were calculated. The results indicate that CoVaccine HT™ enhanced the induction of Th1 responses as evidenced by the significantly higher ratios of IgG2a/IgG2b versus IgG1 obtained from groups given either SdTM or SdTM2P with adjuvant compared to those without adjuvant (Fig. 3B, 3C).

**Fig. 3.**
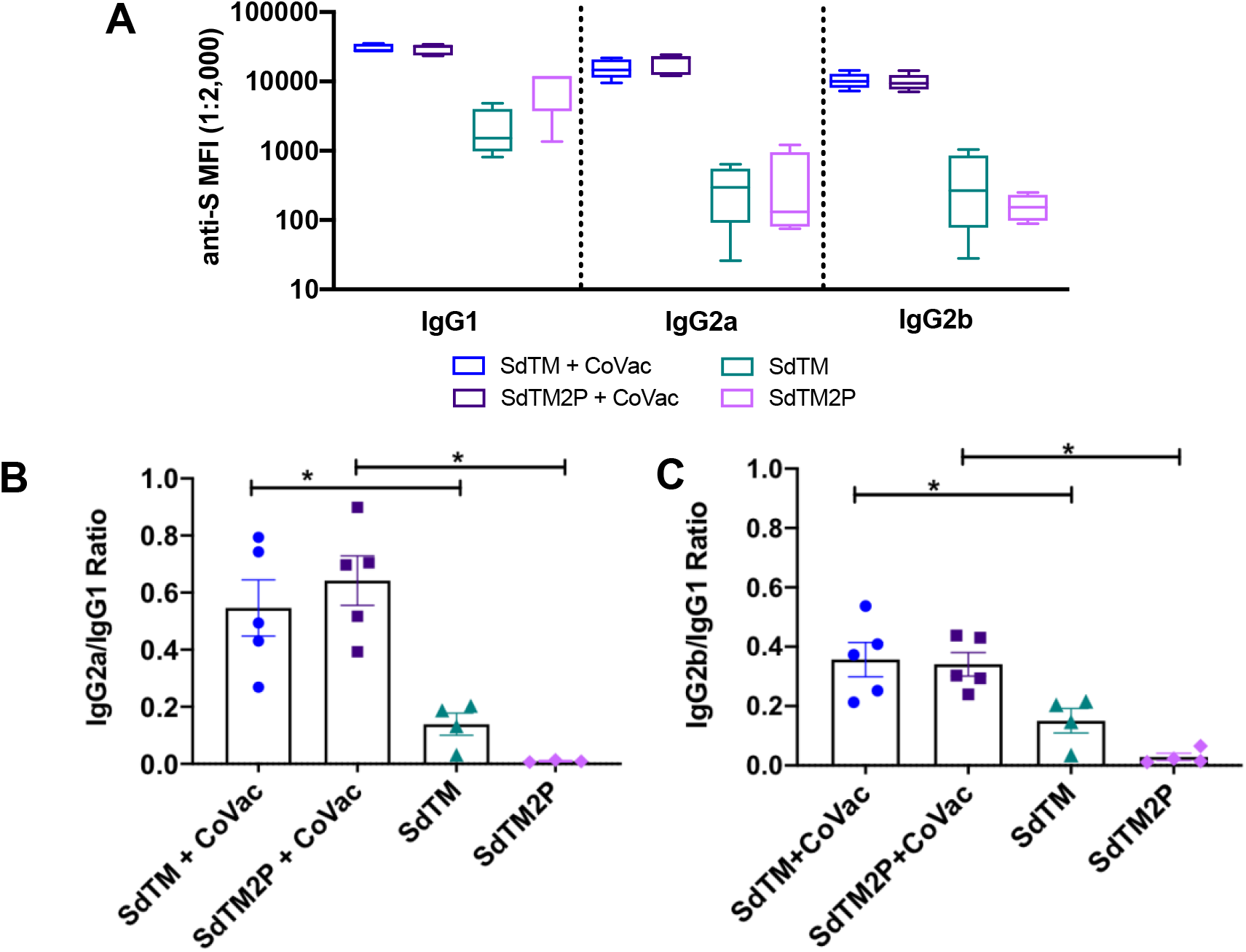
IgG subclass profile induced after vaccination with recombinant S proteins with or without CoVaccine HT™ (CoVac) adjuvant. (A) Mouse sera collected at two weeks post-booster (day 35) were assessed for anti-S specific IgG1, IgG2a, and IgG2b by MIA. The ratios of IgG2a to IgG1 (B) or IgG2b to IgG1 (C) from individual animals were calculated using the MFI values at the serum dilution of 1:2,000. Bars represent mean ± SEM of each group. Significant difference in the ratios between adjuvanted and protein alone groups was determined by Mann-Whitney t test (*p < 0.05).

### 3.4 CoVaccine HT™ adjuvanted SARS-CoV-2 recombinant S protein vaccine elicits IFN-γ T cell response

To assess vaccine-induced T cell responses, we analyzed the number of IFN-γ secreting cells after *ex vivo* stimulation of splenocytes with SARS-CoV-2 spike peptides by a FluoroSpot assay. Stimulation of cells prepared from mice receiving 2 doses of SdTM or SdTM2P with CoVaccine HT™ resulted in robust production of IFN-γ secreting cells (Fig. 4A). Interestingly, one dose of CoVaccine HT™ -adjuvanted S proteins also elicited a rapid IFN-γ secreting response (Fig. 4A). Additionally, a slight decrease in the numbers of IFN-γ secreting cells was observed when mice received a lower amount (2.5 or 1.25 μg) of adjuvanted SdTM2P proteins or a lower amount (0.3 mg) of CoVaccine HT™ (Fig. 4B). However, these differences were not statistically significant. Altogether, our platform shows great potential towards achieving a rapid, robust, and Th1-focused T cell response even in outbred populations.

**Fig. 4.**
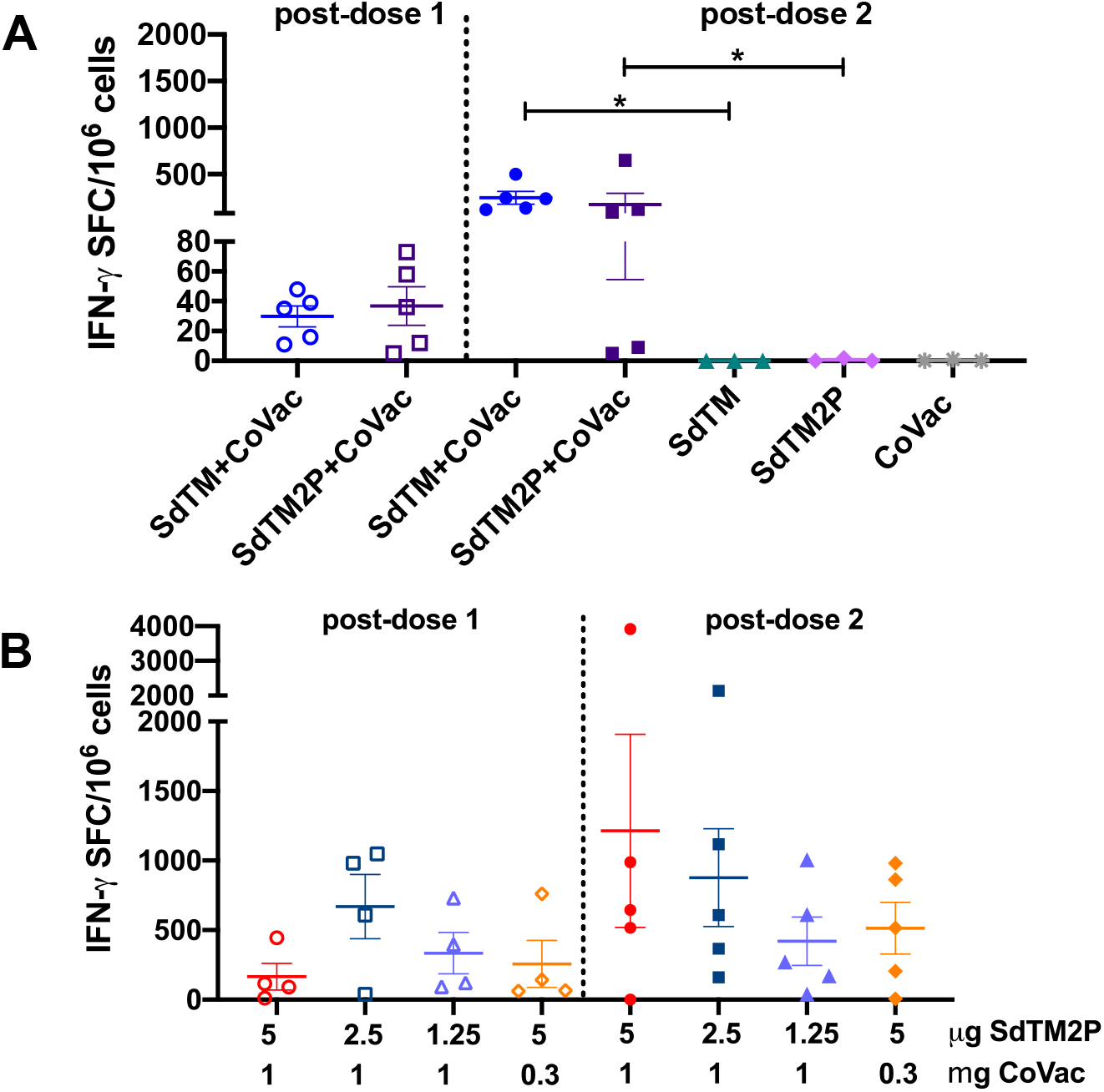
Detection of IFN-γ secreting cells from mice immunized with SARS-CoV-2 vaccines. The splenocytes were obtained from mice (n=3 or 5 per group) immunized with one dose (open symbols) or two doses (closed symbols) of (A) either 5 μg of SARS-CoV-2 SdTM or SdTM2P proteins (purified by hACE2 AC) adjuvanted with 1 mg of CoVaccine HT™ (CoVac) or (B) different dosages (5, 2.5, or 1.25 μ;g) of SdTM2P (purified by mAb IAC) with CoVac (1 or 0.3 mg). The cells were incubated for 24 hours with medium or a peptide pool covering the S protein of SARS-CoV-2 (10 μ;g/mL), and the IFN-γ secreting cells were enumerated by FluoroSpot. The data are shown as the mean values of triplicate assays from individual animals using the number of spot forming cells (SFC) per 10^6^ splenocytes after subtraction of the number of spots formed by cells in medium control wells (< 5 spots). Bars represent mean ± SEM of each group. Significant differences in numbers of IFN-γ secreting cells between groups receiving 2 doses of vaccines with or without adjuvant in panel A was determined by Mann-Whitney t test (*p < 0.05). There are no significant differences between groups given either one or two doses of vaccines when analyzed by one-way ANOVA with Tukey’s multiple comparisons test in panel B.

## Discussion

An ideal COVID-19 vaccine is expected to induce both humoral and cellular immunity, high titers of neutralizing antibodies and Th1-biased response to reduce potential risk of vaccine-associated enhancement of disease [5, 40, 41]. Using a well-established insect cell expression system, we have generated two versions of the SARS-CoV-2 S protein ectodomain and formulated them with a potent adjuvant, CoVaccine HT™. The vaccine candidates elicit both neutralizing antibody and cellular immunity with a balanced Th1/Th2 response in an outbred mouse model. The results obtained in this model may thus inform future vaccine development in animal models more closely related to humans [42]. This supports further preclinical and clinical development of CoVaccine HT™ adjuvanted SARS-CoV-2 vaccine to mitigate the ongoing COVID-19 pandemic.

The S protein of SARS-CoV-2 contains a total of 1,273 amino acids and two major domains (S1 and S2) with distinct structures and functions. Previous preclinical studies of SARS-CoV and MERS-CoV vaccines have demonstrated that the S protein plays a key role in induction of neutralizing antibody and T cell responses as well as protective immunity [16–18]. Stabilization of S proteins in the prefusion trimeric conformation results in increased expression, conformational homogeneity, and production of potent neutralizing antibody responses [18, 27]. Current SARS-CoV-2 vaccines under development use either RBD or full-length S protein with or without modifications for stabilization of prefusion conformation as the major antigen targets. Although RBD is a primary target targeted for potent neutralizing antibodies, it lacks other neutralizing epitopes present on full-length S. This might suggest that full-length S-based vaccines would broaden the neutralizing repertoire and reduce the potential of viral escape from host immunity. In the current work, we produced trimeric S ectodomains in which the furin cleavage site was mutated (SdTM) and that were further stabilized in the prefusion S form by removing the S2’ protease cleavage site and introducing two proline substitutions (SdTM2P). The introduction of two prolines in the S2 subunit resulted in significantly greater production yield (~3-fold increase) in *Drosophila* S2 cells (unpublished data). Vaccination with adjuvanted SdTM or SdTM2P elicits comparable levels of IgG antibody and IFN-γ cellular responses; however, the SdTM2P vaccine generated a slightly higher level of neutralizing antibodies than SdTM, which was also reported in studies of SARS-CoV, MERS-CoV, and other SARS-CoV-2 vaccines [18, 24]. Although more investigation is required to understand whether stabilization of prefusion S ectodomain enhances immunogenicity, the production of stabilized prefusion antigens represents a promising strategy for COVID-19 vaccine design.

CoVaccine HT™ is a novel adjuvant that consists of a sucrose fatty acid sulfate ester (SFASE) immobilized on the oil droplets of a submicrometer emulsion of squalane in water (oil-in-water emulsion) [43]. It has been used for influenza virus and malaria vaccines and shown to enhance humoral and cellular protective immunity, in particular antibody response [44–48]. In addition, we have successfully utilized CoVaccine HT™ in our previous recombinant subunit EBOV, ZIKV and preliminary SARS-CoV-2 vaccine studies and have shown this adjuvant to elicit robust antibody responses [8–10, 29]. Use of CoVaccine HT™ with SdTM and SdTM2P yielded significantly enhanced total IgG and neutralizing antibody responses after both the first and second dose as compared to protein alone which reached similar IgG concentrations after two doses as a single dose of the adjuvanted formulations. Antibody levels (total IgG or neutralizing) did not decrease with a decreased dose of adjuvant, instead a pronounced increase in neutralizing antibody titers were observed, indicating further opportunity for formulation optimization. In addition to improved kinetics, the addition of CoVaccine HT™ modulated the humoral response more towards Th1 type relative to protein alone, as indicated by higher levels of IgG2a and IgG2b. Antigen-specific splenocyte restimulation was more variable, but also increased with the use of adjuvant, particularly after two doses, and were not strongly dependent on adjuvant concentration. These robust responses in outbred mice observed with relatively low antigen and adjuvant doses are particularly encouraging in light of our previous work which has demonstrated the thermostabilization of this adjuvant in combination with other glycoprotein antigens (unpublished data). The similarity of the COVID-19 candidate formulation to prior thermostabilization efforts provides the potential for it to also be similarly thermostabilized in a single vial format. This would allow easier vaccine stockpiling and distribution in regions of the world incapable of maintaining cold-chain logistics necessary for transporting and storing vaccines from other platforms.

The urgent need to rapidly vaccinate the human population worldwide both to slow the increase in fatalities and to prevent the emergence of escape mutations sharply highlights the limitations of vaccines requiring stringent cold chains as even under the best conditions, doses are lost due to inadequate storage or handling. The use of protein subunit vaccines not only offers additional vaccines to be used in parallel to rapidly immunize the global population, but also offers an easier opportunity to develop thermostabilized vaccines which would simplify rapid and far-flung delivery. Using recombinant spike protein without the transmembrane domain and with pre-fusion complex stabilization, we have demonstrated robust immune responses with the CoVaccine HT™ adjuvant in an outbred mouse model. This immune response is observed within 7 days of the first dose and peaks within 14 days after the second dose, showing robust humoral and cell mediated immunity. The immune response was potent with as little as 2.5 μg of protein and 0.3 mg of adjuvant, indicating an economical dose-sparing format should be feasible and should include testing of thermostabilized formulations in rodents and other species.

## Credit author contribution statement

Chih-Yun Lai: Conceptualization, Methodology, Validation, Formal analysis, Investigation, Data curation, Writing-original draft, Writing - review & editing, Supervision, Project administration. Albert To: Conceptualization, Methodology, Validation, Investigation, Resources, Data curation, Writing-original draft, Writing - review & editing, Teri Ann S. Wong: Conceptualization, Methodology, Validation, Investigation, Data curation, Writing - review & editing. Michael M. Lieberman: Conceptualization, Methodology, Formal analysis, Writing - review & editing. David E. Clements: Methodology, Resources, Writing - review & editing. James T. Senda: Methodology, Resources, Writing - review & editing. Aquena H. Ball: Investigation, Writing-original draft. Laurent Pessaint: Investigation. Hanne Andersen: Investigation. Oreola Donini: Writing-original draft, Writing - review & editing. Axel T. Lehrer: Conceptualization, Methodology, Validation, Resources, Writing - review & editing, Visualization, Supervision, Project administration, Funding acquisition.

## Acknowledgments

The authors thank Dr. Andrea Marzi at Laboratory of Virology, National Institute of Allergy and Infectious Diseases for providing rVSV-ΔG expressing SARS-CoV-2 S glycoprotein, Protherics Medicines Development Ltd (London, UK.) for the gift of CoVaccine HT™ adjuvant, and Mapp Biopharmaceutical for the monoclonal antibody CR3022. The authors also acknowledge the following reagent was obtained through BEI Resources, National Institute of Allergy and Infectious Diseases (NIAID), National Institute of Health (NIH): Peptide Array, SARS-Related Coronavirus 2 Spike Glycoprotein, NR-52402.

